## Supplementary for "*sDarken*: Next generation genetically encoded fluorescent sensors for serotonin"

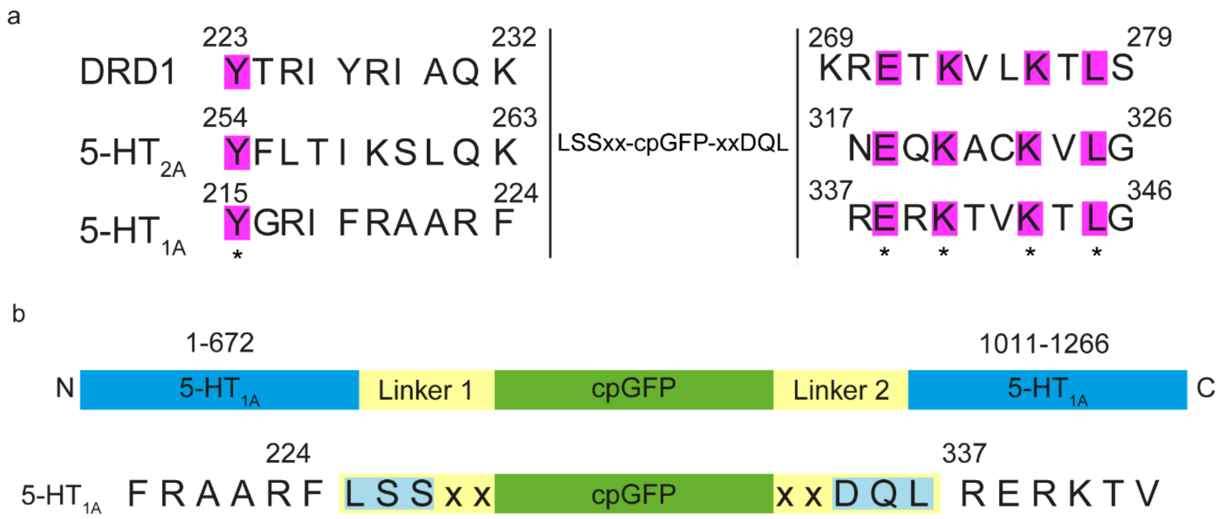


**Supplementary figure 1:** Design principle of *sDarken*. a) the third intracellular loop of the 5-HT1A receptor is replaced by cpGFP. Optimization occurred by mutation of linker 1 and 2. In linker 1 amino acids 4 (position 1) and 5 (position 2) are randomly mutated, whereas in linker two amino acids 1 (position 3) and 2 (position 4 have) been mutated. Amino acid abbreviations: A Ala,C Cys, D Asp, E Glu, F Phe, I Ile, K Lys, L Leu, N Asn, Q Gln, R Arg, S Ser, T Thr, V Val, Y Tyr. b) Sequence alignment of DRD1, 5-HT2A and 5-HT1A

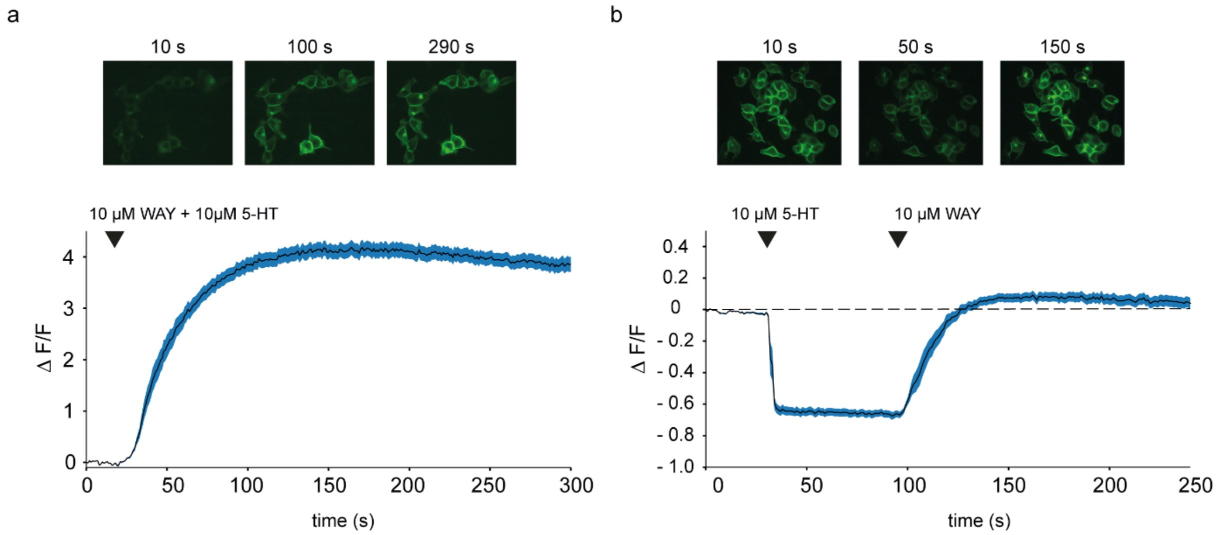


**Supplementary figure 2: Response of *sDarken* to the selective 5-HT$_{1A}$ antagonist WAY 100635.**
a) Wash in of 10 µM WAY and 10 µM 5-HT at frame 15-60. n=24. Bath solution contained 10µM 5-HT.
b) Application of 10 µM 5-HT followed by wash in of 10 µM WAY, n=12. mean ± SEM.

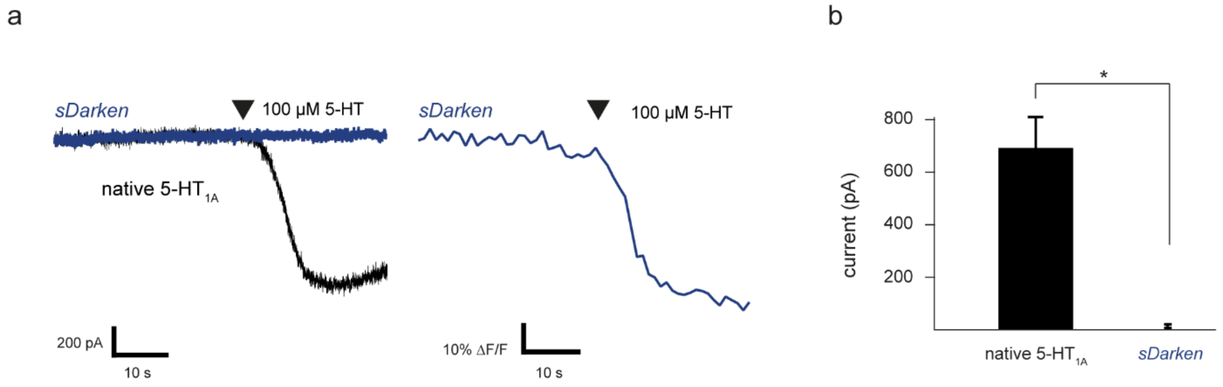


**Supplementary figure 3: *sDarken* has no effect on $G_i$-signaling.** a) Example trace of a GIRK current recording in HEK cells stably expressing GIRK 1 and GIRK 2 subunits. Application of 100 µM 5-HT induced only potassium currents in cells expressing the native 5-$HT_{1A}$ receptor (a, left, black trace), whereas no current response could be observed in HEK cells expressing *sDarken* (a, left, blue trace). b) Comparison of maximal GIRK current amplitude elicited by the application of 100 µM 5-HT (n=3). Values are given as mean ± SEM. *** $p < 0.001$, ** $p < 0.01$. * $p < 0.01$, n.s. not significant

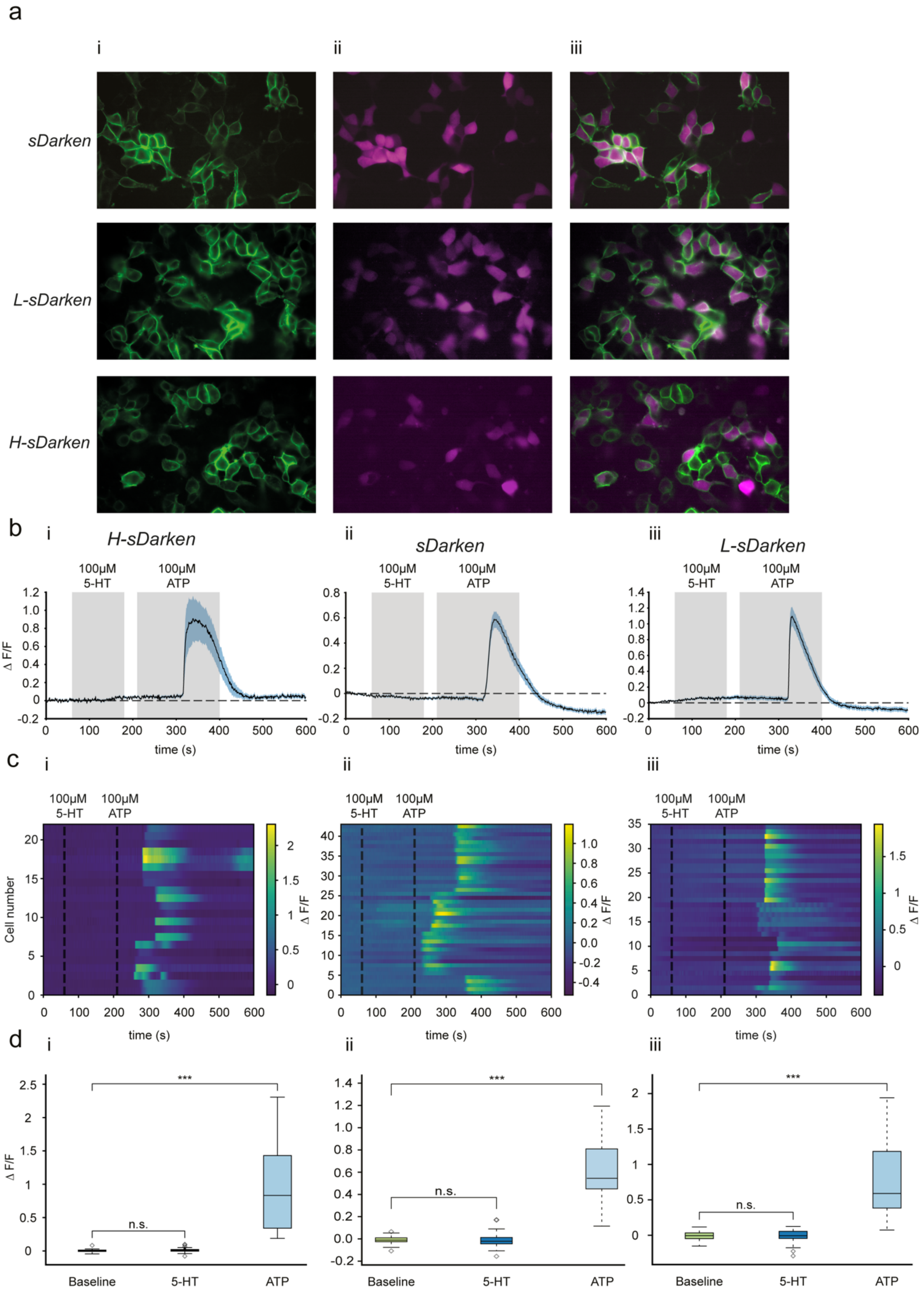
a
i
ii
iii
sDarken
L-sDarken
H-sDarken
b
H-sDarken
sDarken
L-sDarken
100μM 5-HT
100μM ATP
Δ F/F
time (s)
c
Cell number
Δ F/F
time (s)
d
Baseline
5-HT
ATP
n.s.
***

**Supplementary figure 4: s*Darken* variants have no effect on $G_q$-signalling.** (a) Expression of *sDarken*, *L-sDarken* or *H-sDarken* and jRCaMP in HEK cells. i) Fluorescence signal of *sDarken*, *L-sDarken* or *H-sDarken* (green) ii) Fluorescence signal of jRCaMP (magenta). iii) Overlay of *sDarken*, *L-sDarken* or *H-sDarken* and jRCaMP fluorescence signals. (b) Mean ΔF/F values over time (sec) for exemplary trials (i *H-sDarken*: n=6, ii *sDarken*: n=17, iii *L-sDarken*: n=15). Shaded area indicates SEM. Perfusion of cells with 100 µM 5-HT started after 60 seconds and lasted for 120 secs. Subsequently, perfusion of cells with 100 µM ATP started after 220 seconds and lasted for 180 secs. Only cells that co-expressed the 5-HT sensor (*sDarken*, *L-sDarken* or *H-sDarken*) and jRCaMP were included in the analysis. (c) Heatmap of jRCaMP ΔF/F values of all recorded HEK293T cells in all trials (i *H-sDarken*: n=22, ii *sDarken*: n= 43 cells, iii *L-sDarken*: n=35) over time. Dashed lines indicate stimulation of the recorded cells with 100µM 5-HT and subsequently 100 µM ATP. (d) Box-plots of jRCaMP ΔF/F values depicted in (c) before (Timepoint: 10s before 5-HT stimulation), after 100µM 5-HT stimulation (Timepoint: 60s after start of 5-HT stimulation) and after 100µM ATP stimulation (Timepoint: Maximum ΔF/F values after start of ATP stimulation, see Methods). No effect of the 5-HT stimulation on jRCaMP ΔF/F values was observed with any of the three 5-HT sensors (i *H-sDarken*: baseline: -0.0074 ΔF/F ± 0.00573 vs. 5-HT 0.00677 ΔF/F ± 0.00831 (median ± SEM); $p>0.05$ one-way ANOVA repeated measurements, Dunnetts's post-hoc test; ii *sDarken*: baseline: -0.0109 ΔF/F ± 0.005 vs. 5-HT -0.0213 ΔF/F ± 0.009 (median ± SEM); $p>0.05$ one-way ANOVA repeated measurements, Dunnetts's post-hoc test, iii *L-sDarken*: baseline: -0.0064 ΔF/F ± 0.00971 vs. 5-HT -0.00743 ΔF/F ± 0.0162 (median ± SEM); $p>0.05$ one-way ANOVA repeated measurements, Dunnetts's post-hoc test). Stimulation with 100µM ATP significantly increased jRCaMP ΔF/F in HEK293T cells co-expressing either of the three 5-HT sensor variants.(i *H-sDarken*: baseline: -0.0074 ΔF/F ± 0.00573 vs. ATP 0.832 ΔF/F ± 0.141 (median ± SEM); $p<0.05$ one-way ANOVA repeated measurements, Dunnetts's post-hoc test; ii *sDarken*: baseline: -0.0109 ΔF/F ± 0.005 vs. ATP 0.545 ΔF/F ± 0.0403 (median ± SEM); $p<0.05$ one-way ANOVA repeated measurements, Dunnetts's post-hoc test; iii *L-sDarken*: baseline: -0.0064 ΔF/F ± 0.00971 vs ATP 0.590 ΔF/F ± 0.0966 (median ± SEM); ; $p<0.05$ one-way ANOVA repeated measurements, Dunnetts's post-hoc test).

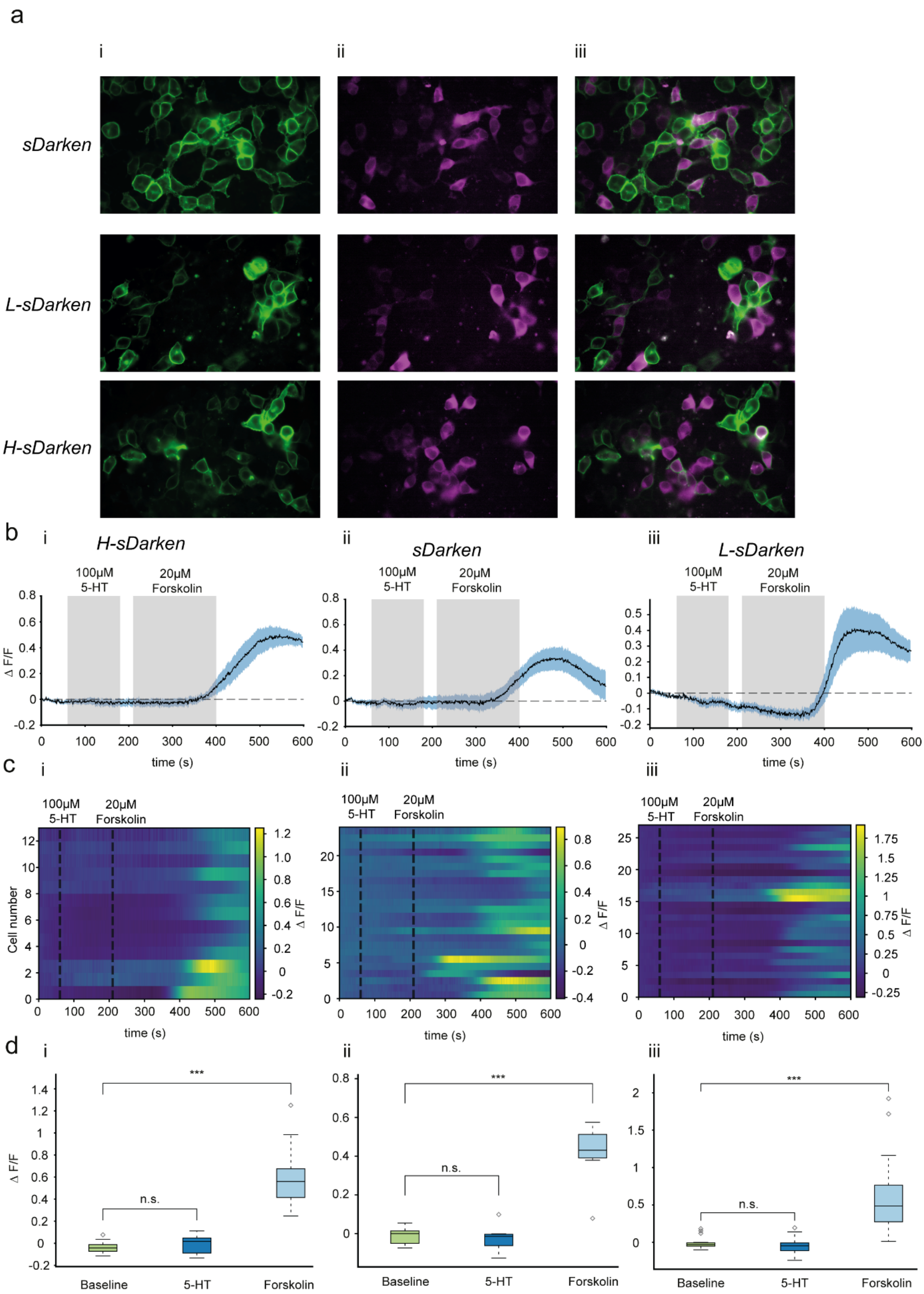
a
i
ii
iii
sDarken
L-sDarken
H-sDarken
b
i
H-sDarken
100μM 5-HT
20μM Forskolin
Δ F/F
time (s)
ii
sDarken
iii
L-sDarken
c
Cell number
time (s)
Δ F/F
d
***
n.s.
Baseline
5-HT
Forskolin

**Supplementary figure 5: Darken variants have no effect om Gs-signalling.** (a) Expression of *sDarken*, *L-sDarken* or *H-sDarken* and CNG-jRCaMP in HEK cells. i) Fluorescence signal of *sDarken, L-sDarken* or *H-sDarken* (green) ii) Fluorescence signal of CNG-jRCaMP (magenta). iii) Overlay of *sDarken*, *L-sDarken* or *H-sDarken* and CNG-jRCaMP fluorescence signals. (b) Mean Delta f/f values over time (sec) for exemplary trials (i *H-sDarken*: n=13, ii *sDarken*: n=5, iii *L-sDarken*: n=4). Shaded area indicates SEM. Perfusion of cells with 100µM 5-HT started after 60 seconds and lasted for 120 secs. Subsequently, perfusion of cells with 20µM Forskolin started after 220 seconds and lasted for 180 secs. Only cells that co-expressed the 5-HT sensor (*sDarken*, *L-sDarken* or *H-sDarken*) and CNG-RCaMP were included in the analysis. **c)** Heatmap of CNG-jRCaMP ΔF/F values of all recorded HEK293T cells in all trials (i *H-sDarken*: n=13 cells, ii *sDarken*: n= 24 cells, iii *L-sDarken*: n=27 cells) over time. Dashed lines indicate stimulation of the recorded cells with 100µM 5-HT and subsequently 20 µM Forskolin. (d) Box-plots of CNG-jRCaMP ΔF/F values depicted in (c) before (Timepoint: 10s before 5-HT stimulation), after 100µM 5-HT stimulation (Timepoint: 60s after start of 5-HT stimulation) and after 20µM Forskolin stimulation (Timepoint: Maximum ΔF/F values after start of Forskolin stimulation, see Methods). No effect of the 5-HT stimulation on CNG-jRCaMP ΔF/F values was observed with any of the three 5-HT sensors (i *H-sDarken*: baseline: -0.00558 ΔF/F ± 0.00592 vs. 5-HT: 0.00785 ΔF/F ± 0.00871 (median ± SEM); $p>0.05$ one-way ANOVA repeated measurements, Dunnetts's post-hoc test; ii *sDarken***:** baseline: baseline: -0.0144 ΔF/F ± -0.0313 vs. 5-HT: -0.0313 ΔF/F ± 0.0153 (median ± SEM); $p>0.05$ one-way ANOVA repeated measurements, Dunnetts's post-hoc test, iii *L-sDarken***:** baseline: -0.0306 ΔF/F ± 0.0128 vs. 5-HT: -0.0469 ΔF/F ± 0.0181 (median ± SEM); $p>0.05$ one-way ANOVA repeated measurements, Dunnetts's post-hoc test). Stimulation with 20µM Forskolin significantly increased CNG-jRCaMP ΔF/F in HEK293T cells co-expressing either of the three 5-HT sensor variants.(i *H-sDarken***:** baseline: -0.00558 ΔF/F ± 0.00592 vs. Forskolin: 0.852 ΔF/F ± 0.147 (median ± SEM); $p<0.05$ one-way ANOVA repeated measurements, Dunnetts's post-hoc test; ii *sDarken***:** baseline: -0.0144 ΔF/F ± -0.0313 vs. Forskolin: 0.409 ΔF/F ± 0.0429 (median ± SEM); $p<0.05$ one-way ANOVA repeated measurements, Dunnetts's post-hoc test; iii *L-sDarken***:** baseline: -0.0306 ΔF/F ± 0.0128 vs Forskolin 0.484 ΔF/F ± 0.0904 (median ± SEM); ; $p<0.05$ one-way ANOVA repeated measurements, Dunnetts's post-hoc test).

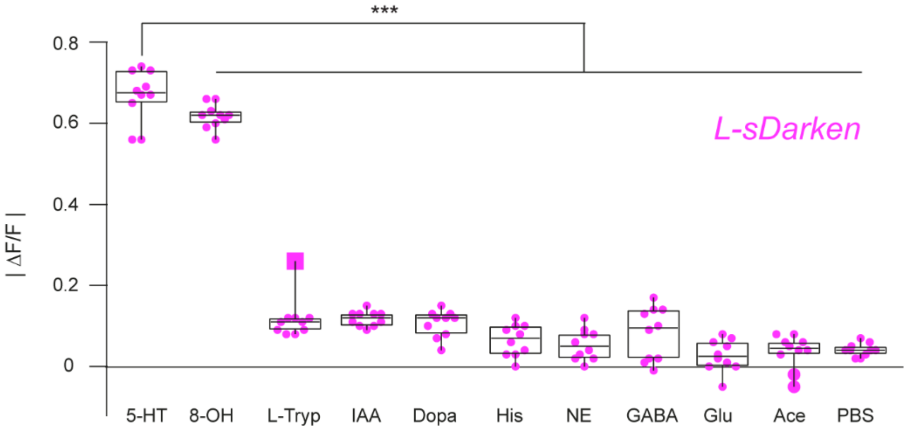


**Supplementary figure 6: Specificity of *L-sDarken* in response to high concentrations**
Fluorescence change to application of serotonin (1.6 mM), 8-OHDPAT (300 µM), similar substances or neurotransmitters if not mentioned differently 3 mM were applied (n=10). Values are given as mean ± SEM. *** $p < 0.001$, ** $p < 0.01$. * $p < 0.01$, n.s. not significant

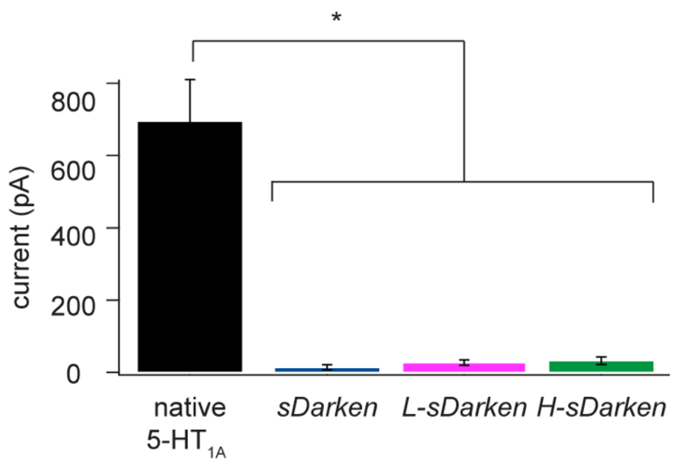


**Supplementary figure 7: GIRK current in response to application of 5-HT.** Application of 100 µM 5-HT induced only potassium currents in cells expressing the native 5-$HT_{1A}$ receptor, whereas no current response could be observed in sensor expressing HEK cells. Comparison of maximal GIRK current amplitude elicited by the application of 100 µM 5-HT or 1 mM 5-HT for *L-sDarken* (5-$HT_{1A}$ n=3, *sDarken* n=3, *L-sDarken* n=4, *H-sDarken* n=4. Values are given as mean ± SEM. *** $p < 0.001$, ** $p < 0.01$. * $p < 0.01$, n.s. not significant

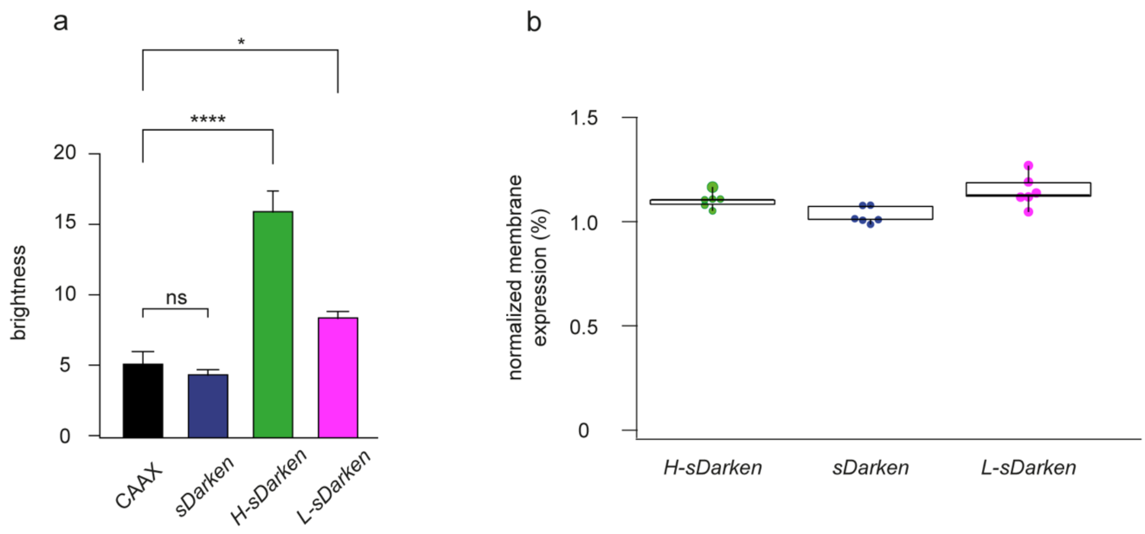


**Supplementary figure 8: Brightness and expression of *sDarken* variants.**
a) Fluorescence intensity of CAAX and the different sensor variants (expressed in HEK cells) after background subtraction (n=20 cells each). Significant differences could be detected between CAAX and *H-sDarken* ($p<0.0001$, Dunnett's multiple comparisons test) and CAAX and *L-sDarken* ($p=0.0352$, Dunnett's multiple comparisons test). b) Membrane expression was calculated as membrane to cytosol fluorescence ratio normalized to CAAX-eGFP (n=6 cells) (Patriarchi et al. 2018).

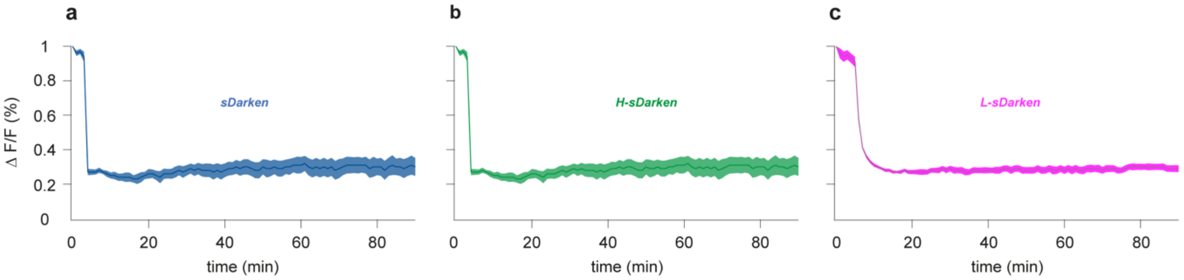


**Supplementary figure 9: Long-term exposure to serotonin.** Fluorescence response of sensor transfected cells to the application of 5-HT for 90 mins. a) *sDarken* application of 100 µM 5-HT, n=5 b) *H-sDarken* application of 100 µM 5-HT, n=5 c) *L-sDarken* application of 1 mM 5-HT, n=5. mean ± SEM

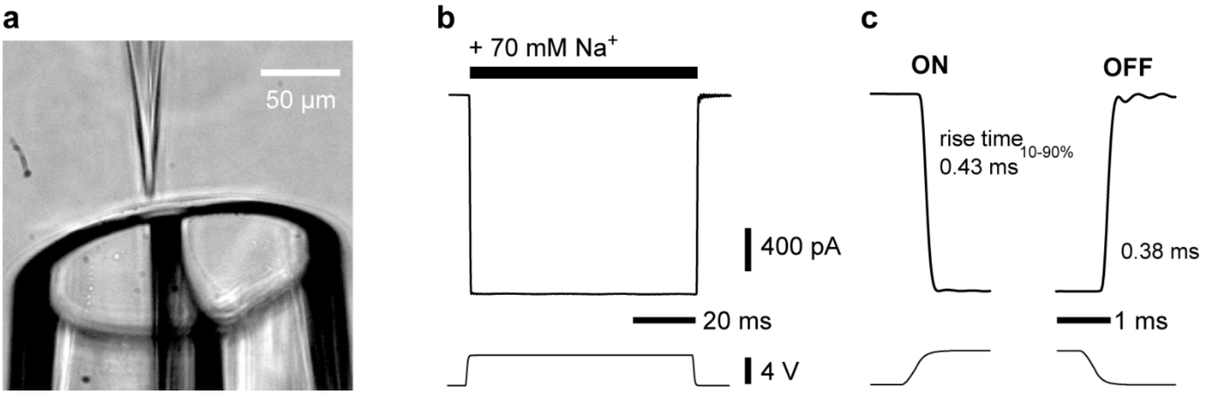


**Supplementary figure 10: Fast piezo-driven solution exchange.** (a) A patch pipette with an excised outside-out patch (top) is placed in front of a piezo-driven double-barreled glass pipette (bottom). One channel contains ligand, the other channel extracellular solution only. For details on the solution exchange system see Pollok & Reiner, 2020. (b) Measurement of fast solution exchange. Exchange currents were monitored by switching between and 70 mM and 140 mM NaCl (downward deflections) at an open patch pipette in voltage-clamp mode. The filtered voltage step, which drives the piezo element, is shown at the bottom. (c) Details of the ON and OFF phase. The current rise and decay times 10-90% are in the submillisecond range. For details see SI Methods.

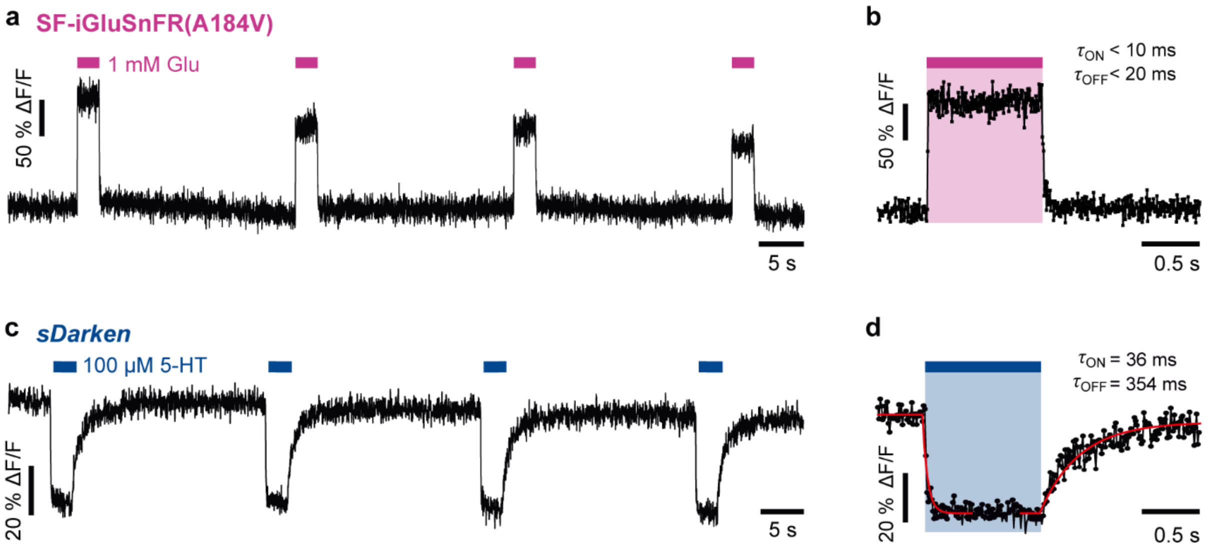


**Supplementary figure 11: Comparison to iGluSnFR.** (a) Repeated application of 1 mM glutamate to a patch from a HEK cell expressing SF-iGluSnFR(A184V) (Marvin et al. 2018) results in fluorescence increases. (b) The example shows a single sweep with ~160% signal change. The ON/OFF kinetics of SF-iGluSnFR(A184V) are fast compared to imaging (194 fps). For the kinetics of iGluSnFR/SF-iGluSnFR see also Helassa et al. 2018 and Marvin et al. 2018. (c) *sDarken* ON/OFF responses upon application of 100 µM 5-HT. (d) Single sweep with ~40% signal change. The kinetics are reasonably well described by single exponential fits (red lines; cf. Fig. 4b) with time constants as indicated. For details see SI Methods.

**a**

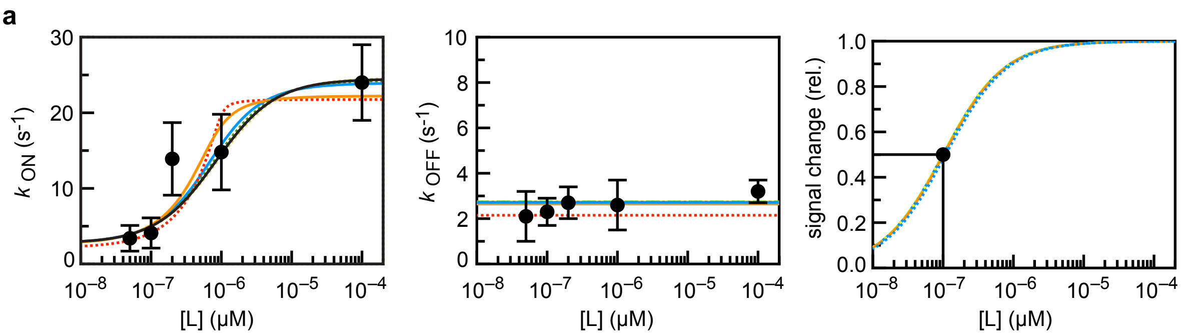


| fifit | $k_{12}$ ($10^7$/(M s)) | $k_{21}$ (1/s) | $k_{23}$ (1/s) | $k_{32}$ (1/s) | chi$^2$ | $k_{12}$ / $k_{21}$ ($10^6$/M) | $k_{OFF}^{app}$ (1/s) | $K_d^{app}$ ($10^{-7}$ M) |
|---|---|---|---|---|---|---|---|---|
| | 2.00 $^{fix}$ | 2.435 | 2.315 | 19.439 | 10.250 | 8.21 | 2.1473 | 1.879 |
| | 3.05 $^{fix}$ | 6.368 | 11.446 | 10.773 | 4.594 | 4.79 | 2.6442 | 1.0122 |
| | **6.109** | **32.540** | **19.439** | **4.488** | **4.122** | **1.88** | **2.7171** | **0.9999** |
| | 12.0 $^{fix}$ | 85.005 | 20.892 | 3.424 | 4.161 | 1.41 | 2.7303 | 0.9999 |
| | 24.0 $^{fix}$ | 192.295 | 21.427 | 3.045 | 4.202 | 1.25 | 2.7353 | 0.9999 |
| | 100.0 $^{fix}$ | 872.389 | 21.769 | 2.807 | 4.239 | 1.15 | 2.7386 | 0.9965 |

**b**

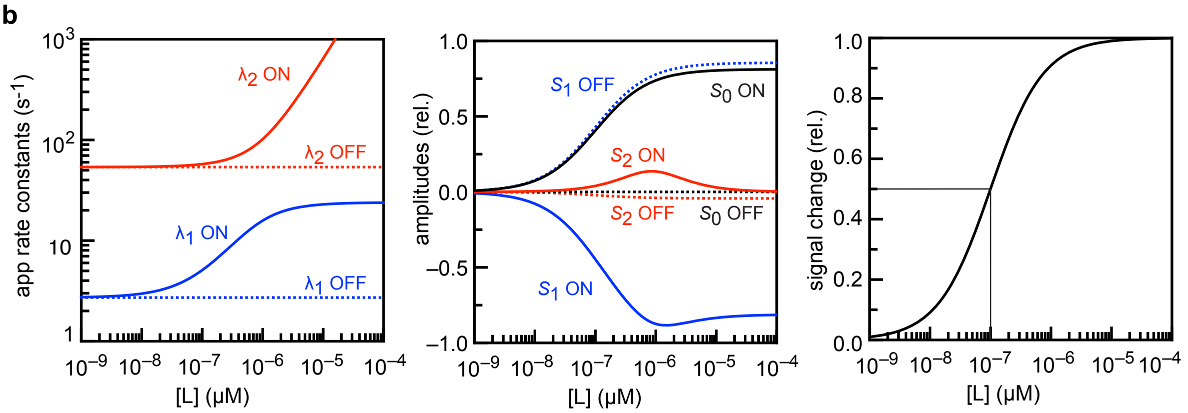


**Supplementary figure 12: *sDarken* signal changes are reproduced by a three-state model**. (a) Fitting of the model (Eq. 1) to the experimental data $k_{ON}$ (*left*), $k_{OFF}$ (*middle*) and $K_D$ (*right*). The freely fitted parameter set ($k_{12} = 6.1 \cdot 10^7$ M$^{-1}$ s$^{-1}$) is shown in blue. Parameters sets obtained by fitting with fixed $k_{12}$ rate constants show similar fit quality, if $k_{12} > 3 \cdot 10^7$ M$^{-1}$ s$^{-1}$ (for details see SI Note Xkineticmodel). (b) Apparent rate constants $\lambda_1$ and $\lambda_2$ (*left*), corresponding amplitudes $S_0$, $S_1$ and $S_2$ (*middle)*, and normalized *sDarken* signal change (*right*) as a function of the ligand concentration [L] calculated for the freely fitted rate constants. Under these conditions, the ON kinetics (solid lines) and OFF kinetics (dashed lines) are dominated by $S_1$, i.e. the experimentally observed signal changes mostly obey single exponential kinetics.

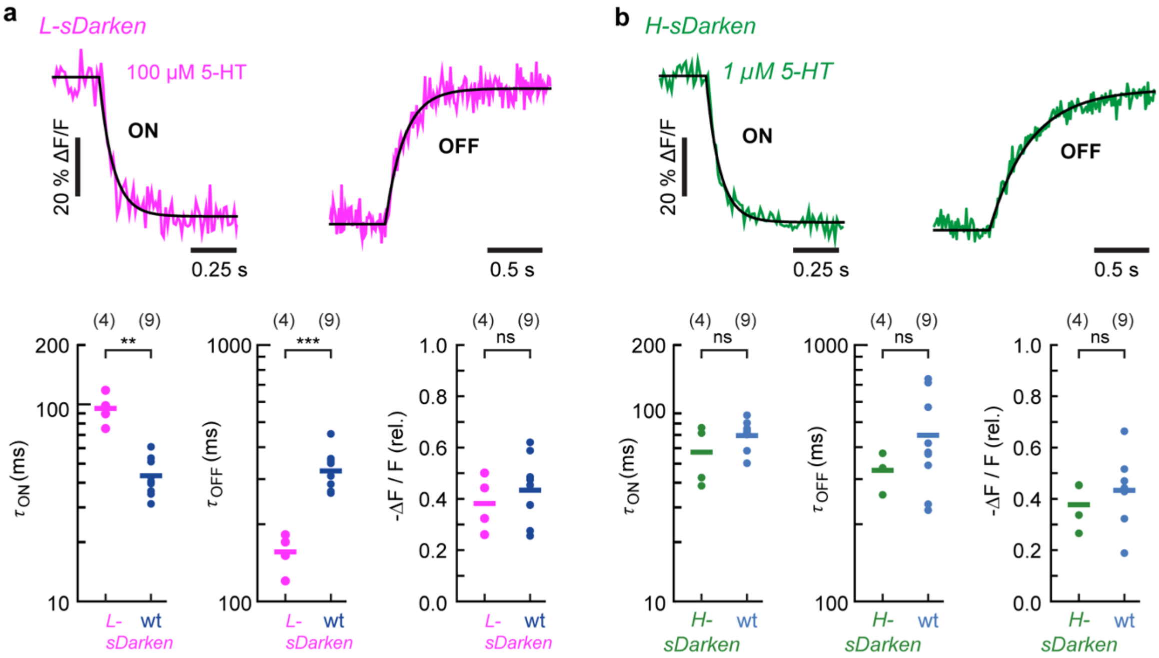


**Supplementary figure 13: ON/OFF kinetics of *L-sDarken* and *H-sDarken*.** (a) ON/OFF kinetics of L-*sDarken* (pink) upon application of 100 µM 5-HT *Top:* Representative trace (average of 4 sweeps) with single exponential fits. *Bottom:* Quantification shows significantly slower ON and significantly faster OFF kinetics compared to s*Darken (wt)* at 100 µM 5-HT, but unaltered signal changes (number of patches given in parenthesis, means shown as crosses). (b) ON/OFF kinetics of *H-sDarken* (green) upon application of 1 µM 5-HT. *Top:* Representative trace (average of 5 sweeps) with single exponential fits. *Bottom:* Quantification shows similar ON kinetics, OFF kinetics and signal changes compared to s*Darken (wt)* at 1 µM 5-HT (number of patches given in parenthesis, means shown as bars). Statistical testing was performed using Welch's *t*-test, ** $p < 0.01$, *** $p < 0.001$. + boxes

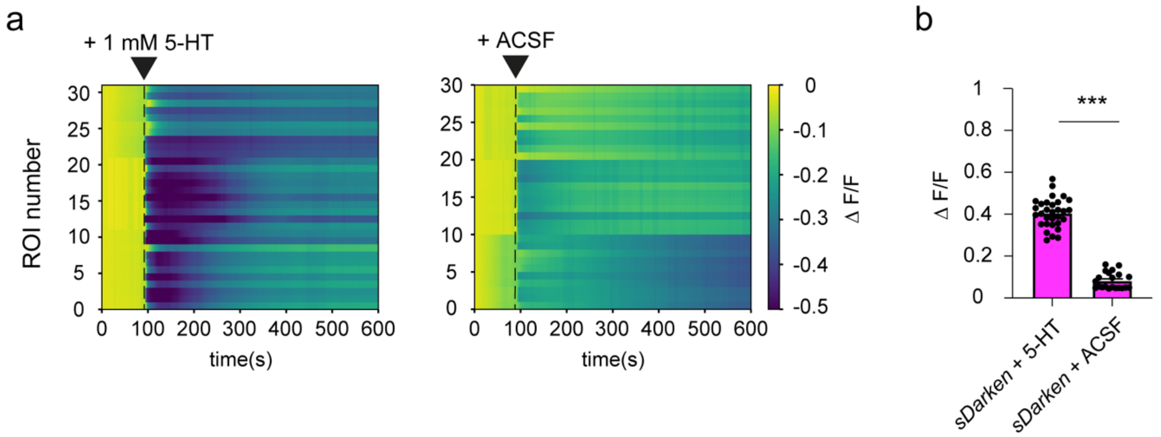


**Supplementary figure 14: Puff application in brain slices, that express *L-sDarken* (a)** Left panel puff application of 1mM 5-HT, n=30 ROIs from 3 slices. Right panel puff application of ACSF, n=30 ROIs in 3 slices. (b) Quantification of fluorescence decrease after puff application from data in a, *L-sDarken* +5-HT 0.41±0.01, *L-sDarken*+ ACSF 0.08±0.008, mean± SEM *** p < 0.001.

### Supplementary Note

*Kinetic model describing the 5-HT sensor*

We investigated the concentration dependence of s*Darken* (**Fig. 4**). First, the observed ON kinetics became faster with increasing 5-HT concentrations, but in the high concentration range (1-100 µM) the ON kinetics did not increase beyond 15-25 $s^{-1}$. This indicates that steps other than 5-HT binding become rate limiting for producing the observed signal change (s*Darken**, fluorescence decrease). The most simple description of the kinetics can thus be given by a three-state scheme (**Fig. 4** and **Eq. 1**), where the 5-HT binding/unbinding equilibrium ($k_{12}$, $k_{21}$) is followed by conformational changes ($k_{23}$, $k_{32}$), which result in a reduced cpGFP fluorescence.

$$\text{L + Darken} \underset{k_{21}}{\overset{k_{12}\cdot[\text{L}]}{\rightleftharpoons}} \text{L•Darken} \underset{k_{32}}{\overset{k_{23}}{\rightleftharpoons}} \text{L•Darken*} \qquad \text{Eq. 1a}$$

$$\text{A} \underset{k_{21}}{\overset{k_{12}'}{\rightleftharpoons}} \text{B} \underset{k_{32}}{\overset{k_{23}}{\rightleftharpoons}} \text{C*} \qquad \text{Eq. 1b}$$

For binding, pseudo-first order reaction kinetics can be assumed ([L] >> [sensor]), i.e. the rate constant $k_{12}'$ is given by $k_{12}\cdot$[L]. This kinetic scheme predicts double exponential signal changes with two apparent rate constants, $\lambda_1$ and $\lambda_2$ (**Eq. 2**).

$$S(t) = S_0 + S_1\cdot\exp(-\lambda_1\cdot t) + S_2\cdot\exp(-\lambda_2\cdot t) \qquad \text{Eq. 2a}$$

The corresponding amplitudes $S_0$, $S_1$ and $S_2$ associated with 'active' sensor configuration (s*Darken**;C) can be obtained analytically for different concentrations, [L], and starting conditions (ON: $A_{t0}$=1, $B_{t0}$=$C_{t0}$=0; OFF: $A_{t0}$=$A_{eq}$, $B_{t0}$=$B_{eq}$, $C_{t0}$=$C_{eq}$) (**Fig. S12**) (Ikai, 1971).

Experimentally single exponential kinetics were observed, which indicates that the second phase ($\lambda_2$, $S_2$) was associated with small amplitudes and/or negative amplitudes with fast kinetics (fast lag phase). This model can reproduce the experimental observations quite well and fitting with $k_{ON}$, $k_{OFF}$ and $K_d = 10^{-7}$ M yields: $k_{12} = 6.1\cdot10^7$ $M^{-1}$ $s^{-1}$, $k_{21} = 32.5$ $s^{-1}$, $k_{23} = 19.4$ $s^{-1}$ and $k_{32} = 4.1$ $s^{-1}$ (**Fig. S12 a**). The apparent rate constants and amplitudes resulting for these values are shown in **Fig. S12 b**. However, it should be noted that these fits remain poorly defined, as $k_{12}$ (and subsequently $k_{21}$) can be increased by order of magnitudes without strongly impacting the quality of neither the fits nor the other parameters (**Fig. S12 a**). Nevertheless, we find that $k_{12}$ is generally $1\text{-}2\cdot10^6$ larger than $k_{21}$ and that reasonable fits are only obtained with $k_{12} > 3\cdot10^7$ $M^{-1}$ $s^{-1}$ (**Fig. S12 a**). In summary, the binding step is fast and already associated with high affinity (0.5-1 µM), whereas the subsequent slower conformational changes appear to be limiting for the observed sensor kinetics.

**Supplementary Movie1. Fluorescence change of *sDarken* in HEK cells** Fluorescence decreases upon application of 800nM 5-HT (single application as indicated in the movie). The movie was acquired with 1 fps and plays in fast forward. Realtime (120 s).

**Supplementary Movie2. Fluorescence change of *sDarken* in HEK cells** Fluorescence decreases upon application of 5µM 5-HT (repetitive application as indicated in the movie). The movie was acquired with 1 fps and plays in fast forward. Realtime (120 s).

**Supplementary Movie3. Fluorescence change of *L-sDarken* in HEK cells.** Fluorescence decreases upon rapid application of 1 µM 5-HT (single application as indicated in the movie). The movie was acquired with 1 fps and plays in fast forward real time (5 s).

**Supplementary Movie4. Fluorescence change of *L-sDarken* in HEK cells.** Fluorescence decreases upon rapid application of 200 µM 5-HT (single application as indicated in the movie). The movie was acquired with 1 fps and plays in fast forward real time (5 s).

**Supplementary Movie 5. Fluorescence change of *L-sDarken* in HEK cells.** Fluorescence decrease upon rapid application of 1600 µM 5-HT (single application as indicated in the movie). The movie was acquired with 91 fps and plays in real time (5 s).

**Supplementary Movie 6. Fluorescence change of *H-sDarken* in HEK cells.** Fluorescence decrease upon rapid application of 50nM 5-HT (single application as indicated in the movie). The movie was acquired with 91 fps and plays in real time (5 s.

**Supplementary Movie 7. Fluorescence change of *H-sDarken* in HEK cells.** Fluorescence decrease upon rapid application of 800nM 5-HT (single application as indicated in the movie). The movie was acquired with 91 fps and plays in real time (5 s).

**Supplementary Movie 8. Mild Bleaching of tddimer2 under continuous two-photon excitation for 1 min.** For details see **Fig. 2**

**Supplementary Movie 9. No bleaching of *sDarken* under continuous two-photon excitation for 1 min.** For details see **Fig. 2**

**Supplementary Movie 10. Fluorescence change of s*Darken* in an outside-out patch.** Fluorescence decrease upon rapid application of 100 µM 5-HT (single application as indicated in the movie). The movie was acquired with 91 fps and plays in real time (5 s). For details see **Fig. 2** and **SI Methods**.

**Supplementary Movie 11: No change in fluorescence of s*Darken* in control condition in organotypic hippocampal slice culture.**

**Supplementary Movie 12:** Change in f**luorescence of s*Darken due to application of 10µM serotonin* in organotypic hippocampal slice culture.**
