## Supplementary material for "*sDarken*: Next generation genetically encoded fluorescent sensors for serotonin": Table 1

| Table 1: Comparison of genetically encoded serotonin sensors | | | | | | | | | |
|---|---|---|---|---|---|---|---|---|---|
| **Sensor** | **($\triangle F/F_0$) % change** | | | | | | **Kd** | **$\tau_{ON}$ (ms)** | **$\tau_{OFF}$= (ms)** |
| | **10 nM** | **100 nM** | **1.6 µM** | **3.2 µM** | **100 µM** | **1 mM** | | | |
| **D116N** | - | - | - | - 20 | - 48 | -73 | 145 µM | 95 | 156 |
| ***Darken*** | - 14 | - 33 | - 58 | - 60 | -40 | Sat. | 127 nM | 43 | 323 |
| **sfDarken** | - 8 | - 51 | - 62 | - 67 | Sat. | Sat. | 57 nM | 57 | 324 |
| **$GRAB_{5\text{-}HT}$**[1] | 70 | 240 | 280 | Sat | Sat | Sat | 22 nM | 200 | 3100 |
| **iSero** [2] | 15 | 20 | 30 | 75 | 750 | Sat | 390 µM | 0.5-10 (fast)<br>5-18 (slow) | 4 (fast)<br>150 (slow) |
| **PsychLight2**[3] | 25 | 55 | 80 | Sat | Sat | Sat | 26.3 nM | | 997 (fast)<br>3998 (slow) |


1. Wan, J. *et al.* A genetically encoded sensor for measuring serotonin dynamics. *Nature Publishing Group* **24,** 746–752 (2021).
2. Unger, E. K. *et al.* Directed Evolution of a Selective and Sensitive Serotonin Sensor via Machine Learning. *Cell* **183,** 1986–2002.e26 (2020).
3. Dong, C. *et al.* Psychedelic-inspired drug discovery using an engineered biosensor. *Cell* **184,** 2779–2792.e18 (2021).
